# ‘Phase Zero’ clinical study platform combining broadband Vis/near-infrared spectroscopy and electrophysiology to study human brain organoid models of neurodevelopmental disorders

**DOI:** 10.1101/2020.06.09.143313

**Authors:** Anirban Dutta, Sneha Sudhakar Karanth, Mahasweta Bhattacharya, Michal Liput, Justyna Augustyniak, Mancheung Cheung, Ewa K. Stachowiak, Michal K. Stachowiak

**Affiliations:** Department of Biomedical Engineering, University at Buffalo, Buffalo, 14260, USA; Department of Pathology and Anatomical Sciences, University at Buffalo, Buffalo, 14260, USA; Polish Academy of Sciences, Mossakowski Medical Research Center, Stem Cell Bioengineering Department, Warsaw, Poland

## Abstract

Homeostatic control of neuronal excitability by modulation of synaptic inhibition (I) and excitation (E) of the principal neurons is important during brain maturation. The fundamental features of in-utero brain developmental, including local synaptic E-I ratio and bioenergetics, can be modeled by cerebral organoids (CO) that have exhibited highly regular nested oscillatory network events. Therefore, we evaluated a ‘Phase Zero’ clinical study platform combining broadband Vis/near-infrared(NIR) spectroscopy and electrophysiology to study E-I ratio based on the spectral exponent of local field potentials and bioenergetics based on the activity of mitochondrial Cytochrome-C Oxidase (CCO). We found a significant effect of the age of the healthy controls iPSC CO from 23 days to 3 months on the CCO activity (χ^*2*^(2,N=10)=20,p=4.5400e-05), and spectral exponent between 30–50Hz (χ^*2*^(2,N=16)=13.88,p=0.001). Also, a significant effect of drugs, choline (CHO), idebenone (IDB), R-alpha-lipoic acid plus acetyl-L-carnitine (LCLA), was found on the CCO activity (χ^*2*^(3,N=10)=25.44,p = 1.2492e-05), spectral exponent between 1–20Hz (χ^*2*^(3,N=16)=43.5,p=1.9273e-09) and 30–50Hz (χ^*2*^(3,N=16)=23.47, p=3.2148e-05) in 34 days old CO from schizophrenia (SCZ) patients iPSC. We present a multidimensional approach combining electrophysiology and Vis-NIR spectroscopy to complement traditional drug design approaches that can advance the system towards a normative parameter space.

## Introduction

Homeostatic control of neuronal excitability by modulation of synaptic inhibition (I) and excitation (E) of the principal neurons during brain maturation is important to avoid runaway excitability conditions^1^. In early brain development, high excitability of the principal neurons may be beneficial to respond to low-input currents^2^; however, the initial phase of postnatal brain development involves a rapid decrease in the E-I ratio with a disproportional increase of inhibitory conductance^1^. Also, prior work^3^ have shown that this rapid neurodevelopment of cortical circuits occurs in an environment of altered energy availability. It can be postulated that the cortical circuit dynamics starts in a reverberating regime with high E-I ratio, and then need to self-organize to different points in the reverberating regime^4^ based on homeostatic plasticity, external inputs^5^, and energy landscape of the developing brain^6^. Here, any runaway excitation or quiescence can lead to abnormal regulation of the E-I balance, which can lead to nervous system disorders, including schizophrenia^7^. In fact, in-utero brain development has been implicated in the pathophysiology of schizophrenia^8^; however, the first symptoms of schizophrenia manifest only in early adulthood^9^. Recent works have shown that brain organoid models can provide an understanding of the fundamental features of in-utero neurodevelopment in health and disease^10^.

Our cerebral organoid studies using induced pluripotent stem cells (iPSCs) from schizophrenia patients and healthy control individuals revealed improperly clustered immature neurons in cortical layers II, III, and V and decreased intracortical connectivity with disrupted orientation and morphology of calretinin interneurons^8^ – a subpopulation of GABAergic cells. Since the GABAergic inhibitory cells locally control the activity of numerous excitatory principal cells, so they maintain local excitation-inhibition (E-I) balance^11^. In fact, E-I balance has been postulated to be an organizing framework for investigating mechanisms in neuropsychiatric disorders^11^, including schizophrenia. Here, calretinin interneurons play a crucial role in the generation of synchronous, rhythmic neuronal activity by interacting with other interneurons^12^. The neuronal activity gives rise to transmembrane currents that can be measured in the extracellular medium^13^, where local synaptic E-I balance can be inferred non-invasively from local field potentials (LFPs) at many spatial scales^14^. Besides LFPs, the spiking output of neurons can also be detected in the extracellular medium^13^, which may together serve as an early biomarker in the development of preventive therapies. The power spectrum of LFPs has been reported to scale as the inverse of the frequency^15^, where Gao et al.^14^ showed that E-I changes could be estimated from the power-law exponent (or, slope). Therefore, GABAergic interneuron defects in schizophrenia^8^ have been postulated to increase the power-law exponent and the E-I ratio from an optimal value (E-I balance hypothesis^11^) and reduce the signal-to-noise ratio (SNR) in the neuronal circuits^11^. Low SNR can make information processing less efficient^11^ which is postulated to be reflected in the bioenergetics^16^.

Brain bioenergetics in health and disease is driven by mitochondrial function^17^. Mitochondria occupies about one-tenth of the total gray matter volume and plays an essential role in the synaptic information processing determining the neuronal performance^18^,^19^. Maintenance of the neuronal activity requires maintained energy supplies, including synaptic transmission – the main energy consumer^20^. The activity of mitochondrial Cytochrome-C Oxidase (CCO), an activator of adenosine triphosphate (ATP) synthase – a marker of mitochondrial function^21^, has been found higher in GABAergic neurons than in the surrounding pyramidal cells^22^. In fact, it has been shown that synaptic boutons with higher synaptic activity contained higher levels of respiratory chain protein cytochrome-c (CytC)^18^, which indicated that the synaptic performance needed to be locally matched with bioenergy supply. Also, mitochondria have been found to regulate neuronal energy metabolism^23^, ^24^. CCO (also known as complex IV) is the terminal complex of the mitochondrial respiratory chain. CCO is comprised of 13 different subunits encoded by three mitochondrial genes (COX subunits I, II, and III) and ten nuclear genes (COX subunits IV, Va, Vb, VIa, VIb, VIc, VIIa, VIIb, VIIc, and VIII) for the molecular pathways involved in mitochondrial energy production. Energy cost in the cortical layers depends on the overall state of homeostatic regulation of global firing rates of neurons determined by the dynamic interaction between the principal cells and the interneurons^22^. Therefore, it can be postulated that calretinin interneurons play a crucial role in this homeostatic regulation by the dynamic interplay of interneurons^12^, where dysfunctional calretinin interneurons can lead to increased excitatory transmission (increased E-I) and excitotoxicity leading to delayed pyramidal cell death^25^. An increased E-I ratio is postulated to lead to an increased CCO concentration that can be measured non-invasively using broadband (490–900 nm) spectroscopy in humans^26^,^21^, as well as in the intact cells and tissues in-vitro^27^,^28^ in our proposed ‘Phase Zero’ clinical study platform^29^,^30^. Here, partial least square processing of the Vis-NIR spectra using redox calibration data can be used for the quantification of CCO in the cerebral organoids.

In schizophrenia, prior works^31^,^32^,^33^ have shown alterations of molecular pathways involved in mitochondrial energy production that is in the cortical layers II and V, which highlighted that the metabolic systems could be novel therapeutic targets^16^. Specifically, a defect in mitochondrial oxidative phosphorylation^34^ with decreased activity of electron transport chain (ETC) complexes, including the reduced activity of complex IV, has been observed^33^. However, it remains to be investigated if promoting mitochondrial function, including ETC, would rescue healthy development in the brain susceptible to schizophrenia. Prior work^35^ on dietary phosphatidylcholine (CHO) ^35^ supplementation starting in the second trimester has shown to activate the timely development of cerebral inhibition based on electrophysiological recordings. At the same time, Idebenone (IDB)^36^, and R-alpha-lipoic acid plus acetyl-L-carnitine (LCLA)^37^ are mitochondrial drugs that are proposed to have an effect on the neonatal pathophysiology related to later schizophrenia risk. In this feasibility study, we investigated the effects of three drugs, CHO^35^, IDB^36^, and LCLA^37^, on cerebral organoids from schizophrenia patients using our ‘Phase Zero’ clinical study platform.

## Methods

### Protocol for organoid development and drug treatment

Cerebral organoids were generated using two healthy control and two schizophrenic cell lines following previously established protocol^8^. The two healthy control cell lines (23476*C: Female and 3651: Female), and the two schizophrenic cell lines (1835: Female and 1792: Male) have been used in our prior works^8,38^. To generate Embryoid Bodies (EBs), confluent induced Pluripotent Stem Cells (iPSC) were maintained on Vitronectin-coated surfaces in combination with Essential 8 (E8) medium in a feeder-free condition. They were then dissociated using EDTA and plated on a 6-well ultra-low attachment plate (Corning, USA) in E8 medium and maintained in the E8 media for 2-3 days. Then, the media was changed to N2/B27 media with dual SMAD inhibitors (which rapidly increases neural differentiation) exchanged daily for 3-4 days. Then, the EBs were transferred into a 24-well low attachment plate (1-2 EBs per well) and grown in neural induction medium for four days. To generate cerebral organoids, neurosperes/neuroepithelium (with visible bright marginal zones) were embedded in matrigel droplets and transferred to 6cm plates in cerebral organoid media without vitamin A for four days, followed by transfer to an orbital shaker and in cerebral organoid media with vitamin A. After ten days in the shaker (14 days after matrigel embedding), the cerebral organoids were treated with a) Vehicle (control), b) Idebenone (IDB), c) Lipoic acid + Acetyl L-carnitine (LCLA), and d) choline (CHO). Media was changed every 2 days, and then the cerebral organoids at various timepoints of maturity were embedded to the center of 3.5cm plate using matrigel, and the broadband vis/near-infrared spectroscopy and electrophysiological investigation were performed. The time points were 34 days for the schizophrenic cerebral organoids, and 23 days, two months, three months for the healthy controls cerebral organoids.

### Experimental setup for broadband vis/near-infrared spectroscopy and electrophysiological investigation

**Figure 1:**
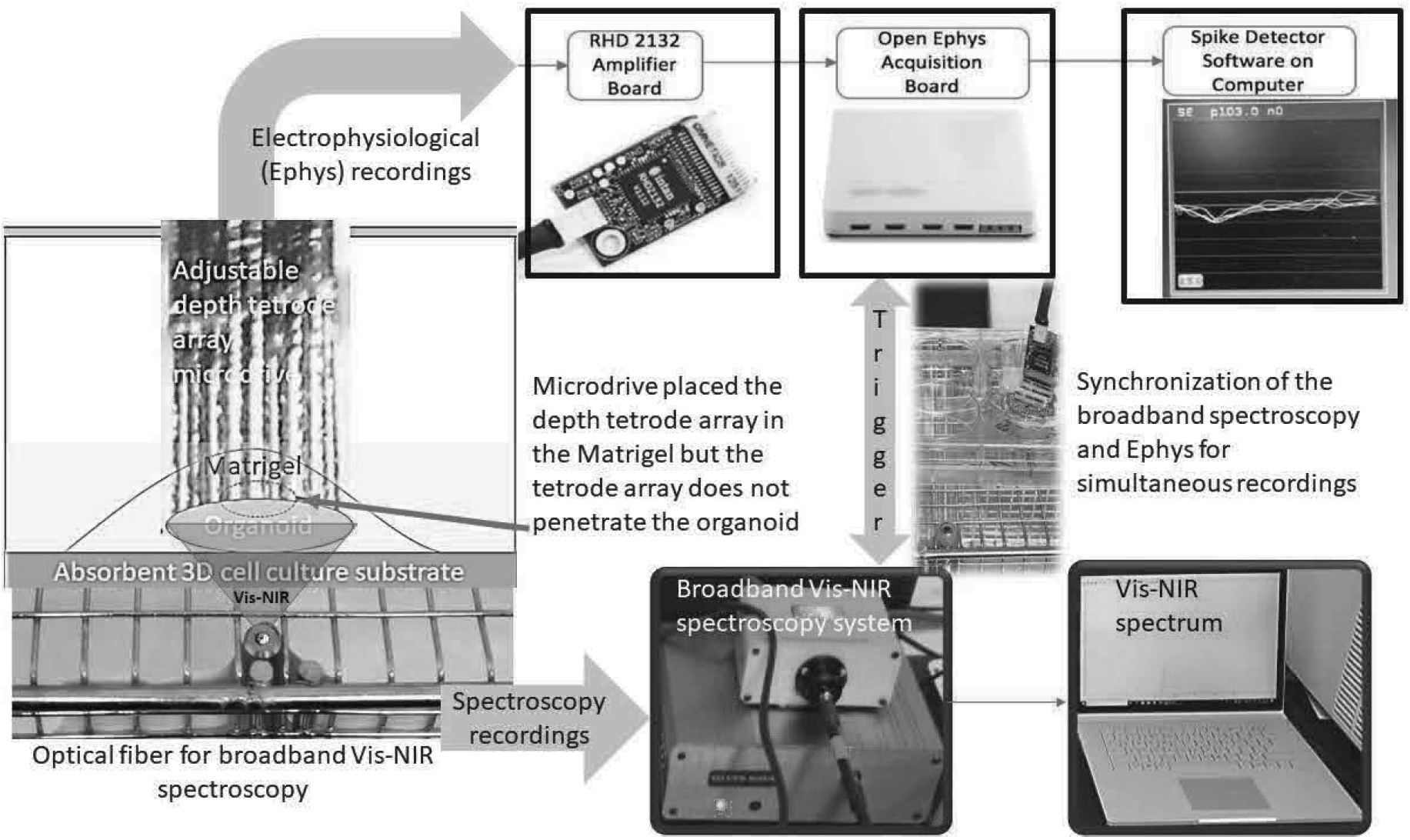
Experimental setup combining broadband vis/near-infrared spectroscopy and electrophysiology. The setup consists of a 32-channel tetrode microdrive integrated with the Intan RHD2132 amplifier (Intan Technologies, USA) and Open Ephys data acquisition system. Intan RHD2132 amplifier (Intan Technologies, USA) supports sampling 32 amplifier channels at 30 kSamples/s each, and provide fully integrated electrophysiology amplifier array with on-chip 16-bit analog-to-digital converter (ADC). We used the Open Ephys GUI and Spike Detector plugin for online monitoring of the data fidelity across the channels as well as for data recording in a laptop computer. Broadband Vis-NIR spectroscopy system consisted of the SILVER-Nova spectrometer (Stellarnet, U.S.A.) over the 190-1110nm wavelength range with 200um slit for high sensitivity, SL1 high stability Tungsten Halogen light (350-2500nm) source (Stellarnet, U.S.A.), and R600-8-VisNIR Reflectance Probe (Stellarnet, U.S.A.) for VIS-NIR spectroscopy with seven optical fibers bundled around one 600μm fiber. We used the SpectraWiz spectrometer software (Stellarnet, U.S.A.) for recording the Broadband Vis-NIR spectroscopy data on a laptop computer.

Figure 1 shows the experimental setup that we developed for broadband vis/near-infrared spectroscopy and electrophysiological investigation. The setup consisted of a 32-channel tetrode (eight polyimide-coated nickel-chrome tetrode wire of diameter ~50μm arranged in a circular grid of ~2mm diameter) microdrive^39^ integrated with Intan RHD2132 amplifier (Intan Technologies, USA) and Open Ephys data acquisition system^40^. The tetrode array was driven into the matrigel matrix^41^ in close proximity to the organoid surface using the microdrive (without penetrating the organoid). The electrical resistivity of the matrigel matrix was found^42^ to be comparable to the skin around 300 ohm.cm in the 100-1000Hz range^43^. Intan RHD2132 amplifier (Intan Technologies, USA) supported sampling 32 amplifier channels at 30 kSamples/sec each and provided a fully integrated electrophysiology amplifier array with on-chip 16-bit analog-to-digital converter (ADC). Open Ephys acquisition board can read from to 8 Intan amplifier chips using low-voltage differential signaling (LVDS) connected to a computer’s USB port^40^. We used the Open Ephys GUI along with the LFP viewer and Spike Detector plugins to record in the fault-tolerant Open Ephys data format in blocks of 1024 samples, each of which included a timestamp and a readily identifiable ‘record marker’^40^. Here, the network events related to the start trigger to synchronize with the broadband Vis-NIR spectroscopy system that was read from the TCP port. Broadband Vis-NIR spectroscopy system consisted of the SILVER-Nova spectrometer (Stellarnet, U.S.A.) over the 190-1110nm wavelength range with 200um slit for high sensitivity, SL1 high stability Tungsten Halogen light (350-2500nm) source (Stellarnet, U.S.A.), and R600-8-VisNIR Reflectance Probe (Stellarnet, U.S.A.) for VIS-NIR spectroscopy with seven optical fibers bundled around one 600μm fiber. We used the SpectraWiz spectrometer software (Stellarnet, U.S.A.) for recording the Broadband Vis-NIR spectroscopy data in conjunction with the electrophysiological recording with Open Ephys GUI.

First, we calibrated the Broadband Vis-NIR spectroscopy system using a colorimetric assay kit (CYTOCOX1, Sigma-Aldrich, USA). We used the broadband (490–900 nm) spectroscopy for the determination of CCO activity (molecular weight 200000 at pH 7^44^) in the standard 2ml sample at various concentrations of 2.5μg/ml, 1.25μg/ml, 0.625μg/ml, 0.3125μg/ml, 0.1562μg/ml, 0.0781μg/ml, 0.05μg/ml, 0.1μg/ml, 0.15μg/ml, 0.2μg/ml, 0.25μg/ml, 0.125 μg/ml, 0.175 μg/ml, as well as in the buffer solution. Principal component regression and partial least square processing of the Vis-NIR spectra were investigated to develop a multivariate regression model for the determination of the CCO activity in the organoid – see Appendix 1. We recorded spontaneous neuronal activity for a total of 10 minutes from 32-channel tetrode along with Vis-NIR spectroscopy in a staggered manner over ten trials each. The spontaneous neuronal activity is a combination of multi-unit activity (MUA) and LFPs that were saved by the Open Ephys GUI (Spike Detector plugin). The electrodes that did not record at least five spikes/min were discarded^45^. More than 50% of the electrodes recorded at least five spikes/min, so 16 best electrodes with most spikes/min were used for LFP analysis. LFPs were resampled at 500Hz, and then processed by power spectral density (PSD) estimates computed using Welch’s method with a window length of 2sec and overlap of 1sec^45^. Then, the spectral exponent^46^ of non-oscillatory PSD background was computed (fitPowerLaw3steps.m) from the PSD under the assumption that LFPs decays according to an inverse power-law: PSD(f) ~1/f^α^. Here, the spectral exponent β = −α indexes the steepness of the decay of the PSD background, typically ranging from −4 to −1.5^47^. It is important to take the non-oscillatory PSD background for the power-law fitting (fitPowerLaw3steps.m) since a forward LFP computational model showed that the oscillatory peaks in the PSD could reduce the correlation of the power-law exponent with the E-I ratio^14^.

In our LFP recordings, both prominent oscillatory peaks, as well as aperiodic signal characteristic, have been observed^45^ – see Appendix 2. Therefore, a three-step fitting procedure^46^ discarded the oscillatory peaks prior to estimating the background slope where fitting was done in 5Hz windows. The average slope in the 30-50Hz frequency range is postulated to be positively correlated with the synaptic E-I ratio based on forward LFP computational model^14^. Here, change in the slope at the higher frequency band (>30Hz) can occur separately than the change in the slope at a lower frequency band (<30Hz) based on prior work on the human cortex^48^. So, we also investigated the average slope in the lower frequency band of 1-20Hz as a control since the lowest frequency oscillatory activity was primarily in the 10-15Hz band^14^. We recorded from a 23 day old, two months old, and three months old cerebral organoids from healthy controls, and a 34 day old cerebral organoid from schizophrenia patients where it was postulated that older organoids from healthy controls would have a higher ratio of inhibitory neurons^45^ (i.e., lower E-I ratio and lower slope). Also, we investigated drug (CHO, IDB, LCLA) treated 34 days old cerebral organoids from schizophrenia patients, which was postulated to have a higher ratio of inhibitory neurons^45^ (i.e., lower E-I ratio, lower slope) when compared with the vehicle-treated control. We used a nonparametric version of the analysis of variance, nonparametric Friedman’s test, that make only mild assumptions about the data, and are appropriate when the distribution of the data is non-normal.

## Results

### Electrophysiology and broadband vis/near-infrared spectroscopy of the cerebral organoids without drug treatment

Multivariate regression modeling using partial least square regression (PLSR) as well as principal component regression (PCR) found that three components accounted for greater than 99% of the variance in the calibration data – see Figure A1(A) in Appendix 1. Also, ten components accounted for 99.999% of the variance and provided an R-squared (the proportion of the variance in the dependent variable that is predictable from the independent variable) goodness-of-fit of 0.9906 for the PLSR model (compared to R-squared PCR10=0.9707 – see Figure A1(B) in Appendix 1). We used ten component PLSR model (PLS10 – see Figure A1(C) in Appendix 1) for estimating the CCO activity (in nM for molecular weight 200,000 at pH 7^44^) across ten trials, as shown in Figure 2(A) as violin plot. Violin plot allowed the visualization of the distribution of the data and its probability density where the box plot (with median, interquartile range, upper adjacent value, lower adjacent value) is combined with the probability density placed on each side. Friedman’s test showed a significant (χ^*2*^(2, N=10) = 20, p = 4.5400e-05) effect of the age of the control cerebral organoids on the CCO activity. Here, the CCO activity decreased with increasing maturity of the cerebral organoids from healthy controls. The CCO activity in the 34-day old cerebral organoids from the schizophrenia (SCZ) patients was found to be comparable to the CCO activity in the two months old healthy control. The corresponding average spectral exponents for the 16 best (most spikes per min) tetrode channels are shown as the violin plot in 1 – 20Hz frequency band in Figure 2(B), and 30 – 50Hz frequency band in Figure 2(C), which were calculated from the corresponding PSDs (shown in Appendix 2). Friedman’s test showed an insignificant (χ^*2*^(2, N=16) = 3.13, p = 0.2096) effect of the age of the healthy cerebral organoids on the spectral exponent in 1 – 20Hz frequency band; however, a significant (χ^*2*^(2, N=16) = 13.88, p = 0.001) effect of the age of the healthy cerebral organoids on the spectral exponent in 30 – 50Hz frequency band. Here, the spectral exponent in 30 – 50Hz frequency band decreased with increasing maturity of the cerebral organoids from the healthy controls, which indicated a decreased E-I ratio. Also, the spectral exponents of the vehicle-treated 34-day old cerebral organoids from the SCZ patients was found to be comparable to that of the two months old healthy control.

**Figure 2:**
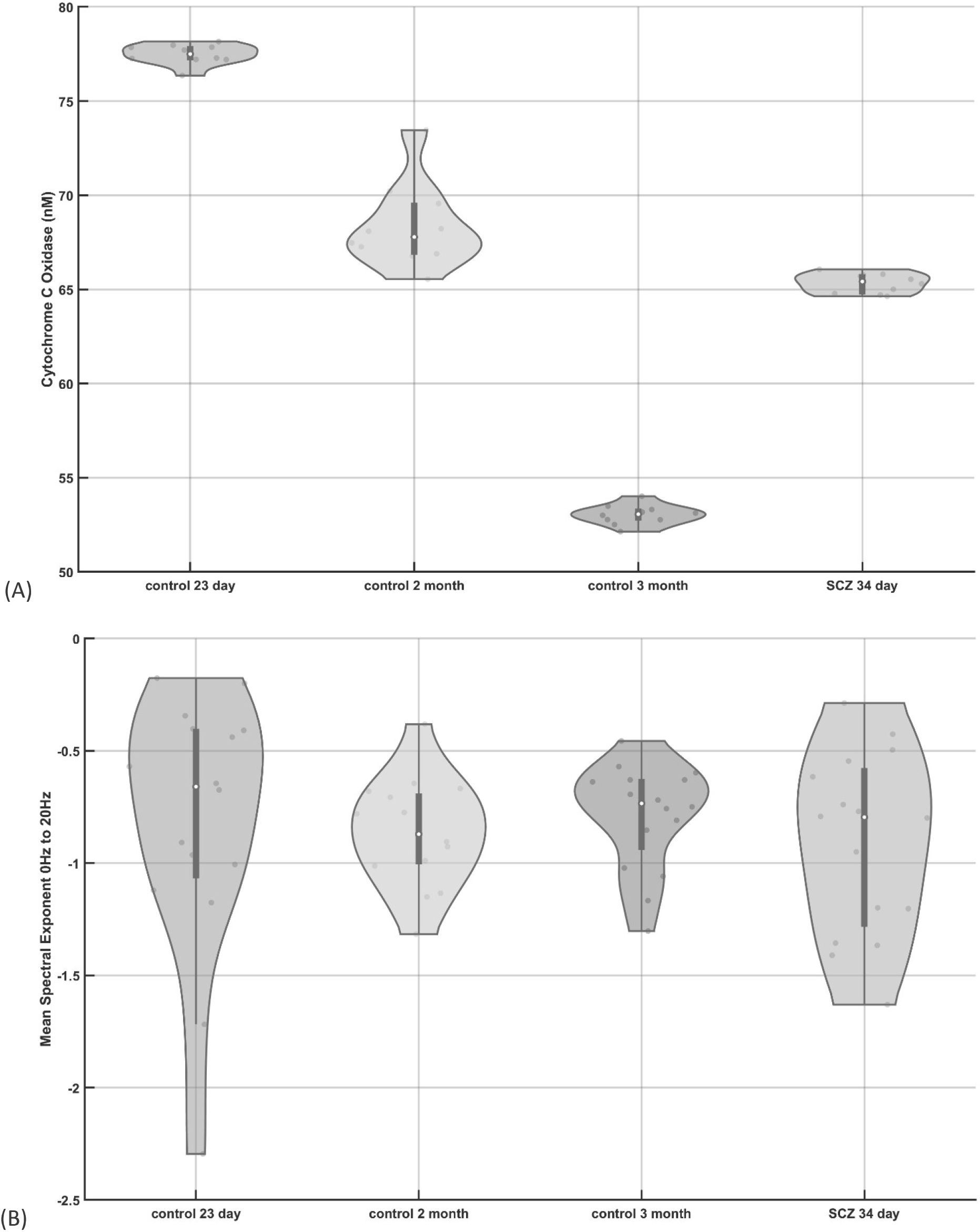
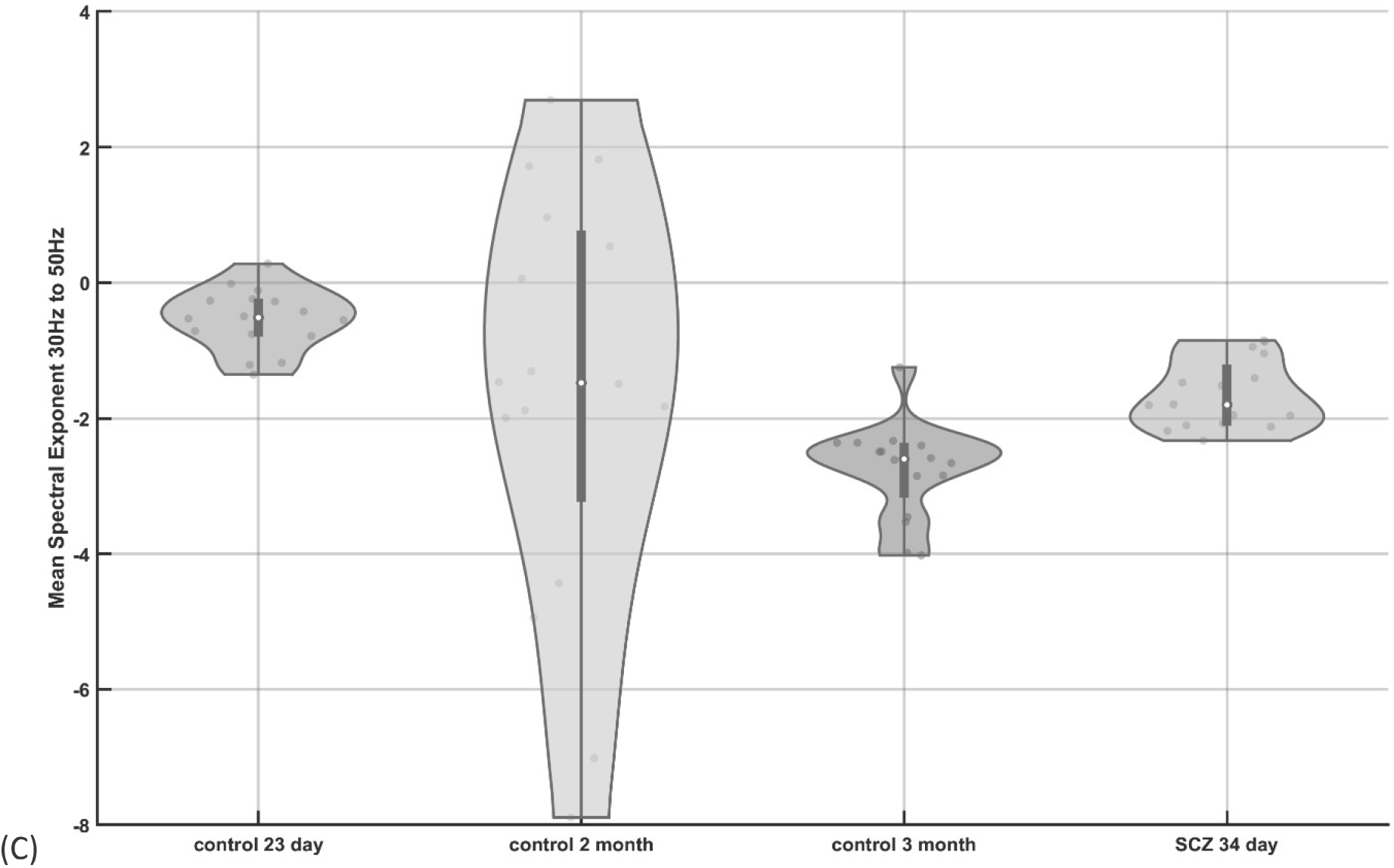
(A) Violin plot of the cytochrome C oxidase (CCO) activity across ten trials in cerebral organoids from healthy controls (control) at 23 days, two months, and three months, as well as a cerebral organoid from schizophrenia (SCZ) patients at 34 days (vehicle-treated). (B) Violin plot of the spectral exponent in the 1 – 20Hz frequency band computed from 16 best (most spikes per min) tetrode recordings from cerebral organoids at 23 days, two months, and three months, for healthy controls (control) as well as a vehicle-treated cerebral organoid at 34 days from schizophrenia (SCZ) patients. (C) Violin plot of the spectral exponent in the 30 – 50Hz frequency band computed from 16 best (most spikes per min) tetrode recordings from cerebral organoids at 23 days, two months, and three months, for healthy controls (control) as well as a vehicle-treated cerebral organoid at 34 days from schizophrenia (SCZ) patients. Violin plot allowed visualization of the distribution of the data and its probability density where the box plot (with median, interquartile range, upper adjacent value, lower adjacent value) is combined with the probability density placed on each side.

### Electrophysiology and Broadband vis/near-infrared spectroscopy of schizophrenia (SCZ) patients cerebral organoids with drug treatment

We used a ten component PLSR model (PLS10) for estimating the CCO activity (in nM) across ten trials of 34-day old cerebral organoids from the schizophrenia (SCZ) patients with drug and vehicle treatment, as shown in Figure 3(A) as violin plot. Friedman’s test showed a significant (χ^*2*^(3, N=10) = 25.44, p = 1.2492e-05) effect of drugs on the CCO activity for the cerebral organoids from the SCZ patients. Here, the CCO activity decreased in the case of CHO and IDB while increased in the case of LCLA when compared to the vehicle-treated one. The corresponding average spectral exponents for the 16 best (most spikes per min) tetrode channels in 1 – 20Hz frequency band and in the 30 – 50Hz frequency band are shown as the violin plot in Figures 2(B) and 2(C) respectively, which were calculated from the corresponding PSDs (shown in Appendix 2). The oscillatory peak between 10 – 15Hz was found to be decreased by the drugs, especially in the case of IDB, when compared to the healthy controls – see Figure A2(D)-(G) (Appendix A2). Friedman’s test showed a significant (χ^*2*^(3, N=16) = 43.5, p = 1.9273e-09) effect of drugs on the spectral exponent in the 1Hz – 20Hz frequency band for the cerebral organoids from the SCZ patient. Also, Friedman’s test showed a significant (χ^*2*^(3, N=16) = 23.47, p = 3.2148e-05) effect of drugs on the spectral exponent in 30 – 50Hz frequency band; however, the effects were different from the 1Hz – 20Hz frequency band. While the spectral exponent in the 1 – 20Hz frequency band was decreased by IDB and LCLA, the spectral exponent increased in the 30 – 50Hz frequency band when compared to the vehicle-treated one. CHO slightly increased the spectral exponent in both the frequency bands when compared to the vehicle-treated one.

**Figure 3:**
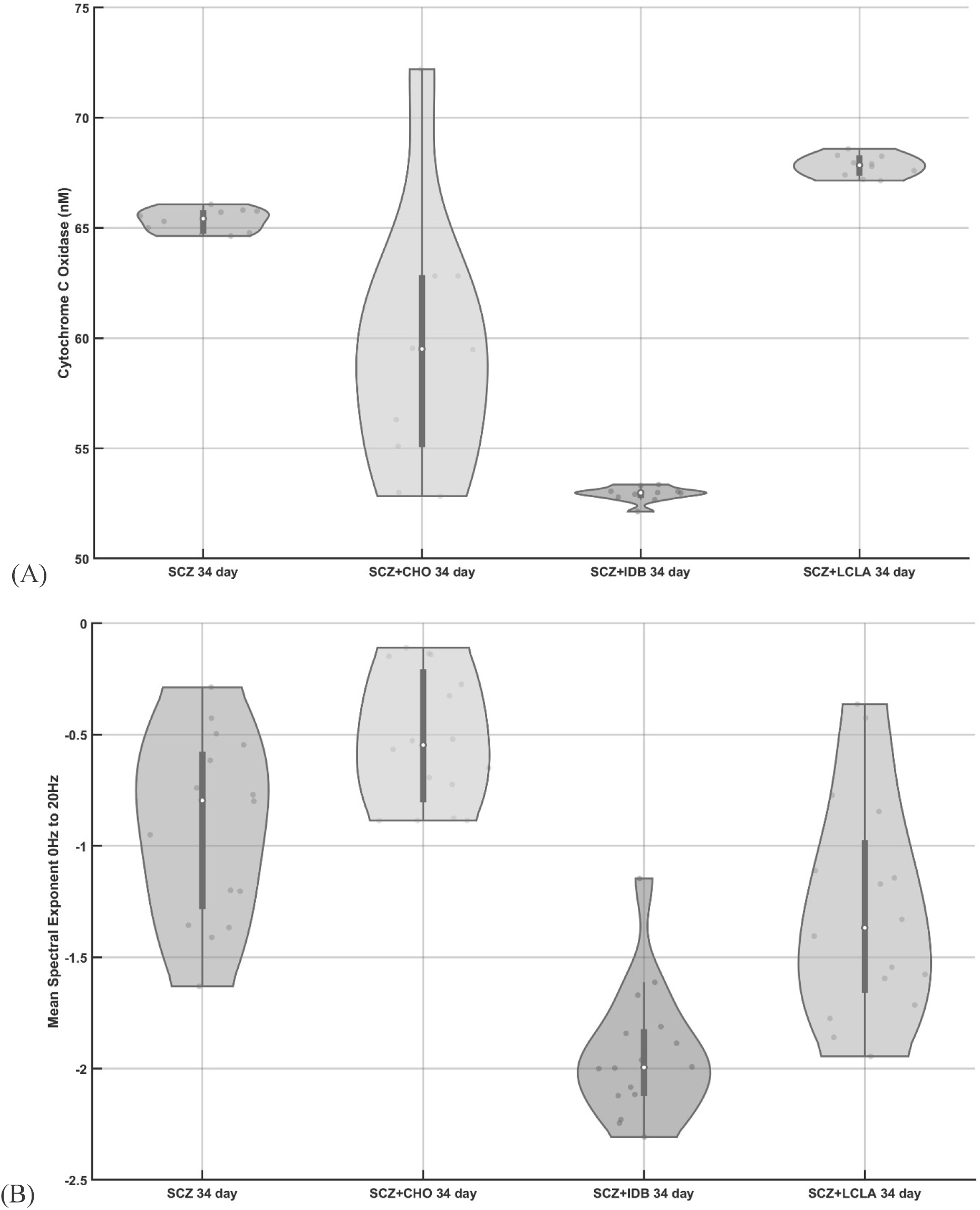
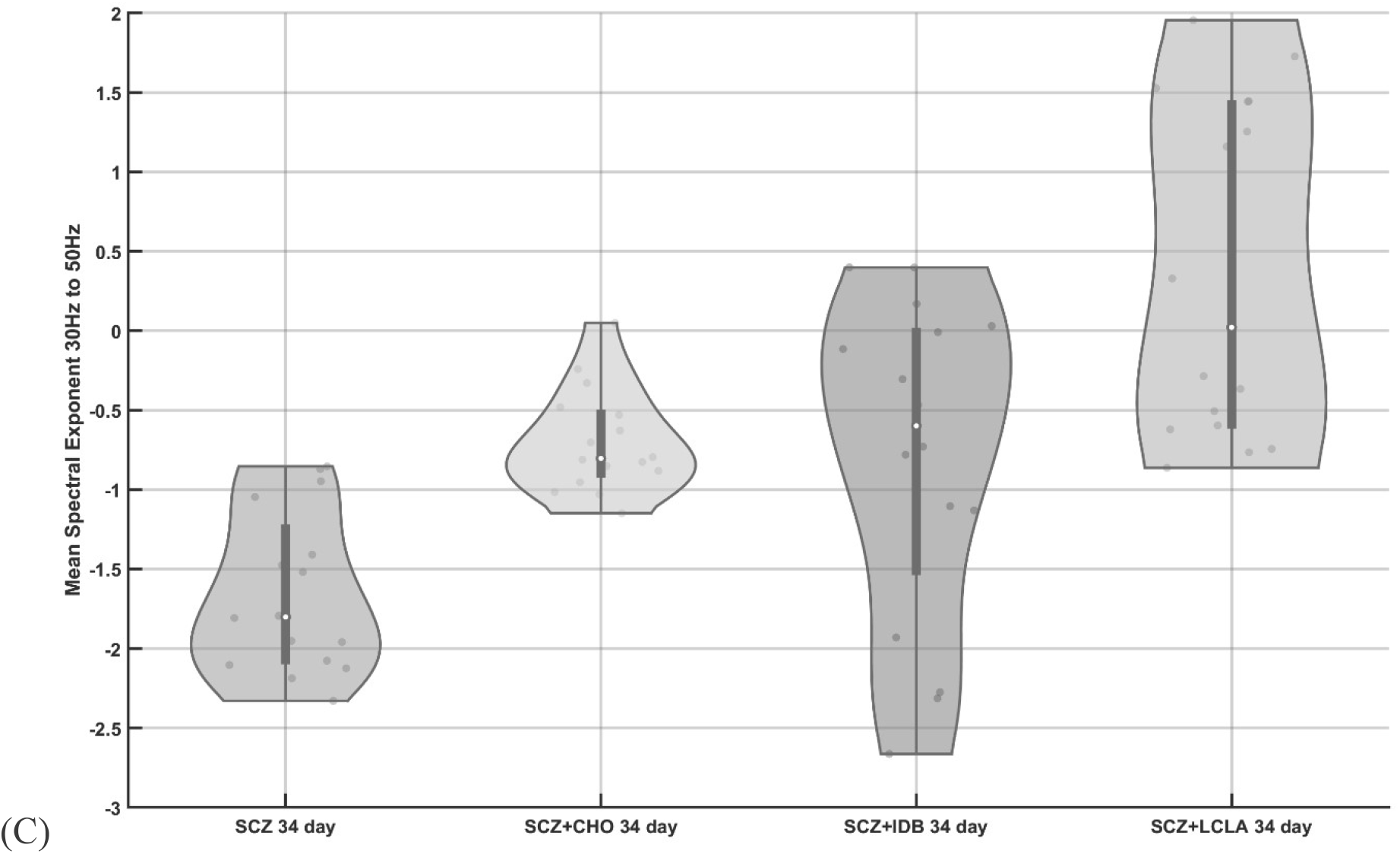
(A) Violin plot of the cytochrome C oxidase (CCO) activity across ten trials in 34 days old cerebral organoids from schizophrenia (SCZ) patients treated with vehicle and drugs, choline (CHO), idebenone (IDB), R-alpha-lipoic acid plus acetyl-L-carnitine (LCLA). (B) Violin plot of the spectral exponent in the 1 – 20Hz frequency band computed for 16 best (most spikes per min) tetrode recordings from 34 days old cerebral organoids from schizophrenia (SCZ) patients, treated with vehicle and drugs, CHO, IDB, LCLA. (C) Violin plot of the spectral exponent in the 30 – 50Hz frequency band computed for 16 best (most spikes per min) tetrode recordings from 34 days old cerebral organoids from schizophrenia (SCZ) patients, treated with vehicle and drugs, CHO, IDB, LCLA. Violin plot allowed visualization of the distribution of the data and its probability density where the box plot (with median, interquartile range, upper adjacent value, lower adjacent value) is combined with the probability density placed on each side.

We also investigated the effects of the drugs on the cerebral organoids from healthy controls that are presented in Appendix 3. All the three drugs reduced the CCO activity in the healthy controls; however, IDB and LCLA had opposite effects on the spectral exponent in the 30 – 50Hz frequency band and no effect on the spectral exponent in the 1 – 20Hz frequency band in the healthy controls when compared to the drug-treated SCZ cerebral organoids at 34 days. CHO had a similar effect on the spectral exponent in the 1 – 20Hz frequency band; however, it increased the spectral exponent in the 30 – 50Hz frequency band in healthy controls when compared to the vehicle-treated condition in the cerebral organoids at 2months. In general, the spectral exponent in the 30 – 50Hz frequency band was most responsive to the drug treatment.

## Discussion

An optimal value of E-I balance has been proposed to be determined by the (near) criticality of the cortical circuit dynamics, which is a regime that maximizes information processing^49^,^50^,^51^,^4^. Cerebral organoids have exhibited periodic and highly regular nested oscillatory network events^45^; however, external inputs^5^ may be necessary to transition to a more in-vivo cortical circuit dynamics based on homeostatic plasticity^52^. In in-vivo cortical networks, Beggs and Plenz^50^ showed that the propagation of spontaneous activity obeys a power law with an exponent of −1.5 for event sizes when close to the critical dynamics. Beggs and Plenz^50^ also presented other works that have shown an exponent around −1.5 to be characteristic of the cortical circuit dynamics. In a computational model, as the E-I ratio was varied from 1:2 to 1:6, the PSD slope of the LFPs between 30 and 50 Hz correlated positively with the E-I ratio^14^. In the current study, the spectral exponent in the 30 – 50Hz frequency band was found (see Figure 2(C)) to decrease with increasing maturity of the cerebral organoids in the healthy controls, which indicated a decrease in the E-I ratio. Also, the CCO activity decreased with increasing maturity of the cerebral organoids from healthy controls. This is postulated to be due to an increased inhibition in the cortical circuit dynamics with increasing maturity of the cerebral organoids where Ma et al.^49^ showed that the criticality in excitatory networks is established by the inhibitory plasticity and architecture. Also, during neurodevelopment, a metabolic shift of cells from glycolysis to oxidative phosphorylation is necessary for an increase in bioenergetics^53^,^54^, and for orchestrating nuclear transcriptional program^55^,^56^. Here, energy supply from mitochondria is essential and can be limiting for synaptic activity during neurodevelopment, which can reciprocally modulate the motility and fusion/fission balance of mitochondria in the dendrites^24^ as well as homeostatic plasticity^57^.

Drugs, IDB and LCLA, had an opposite effect in the two frequency bands – while the spectral exponent was increased in the 30 – 50Hz frequency band, the spectral exponent decreased in the 1 – 20Hz frequency band when compared to vehicle-treatment. CHO slightly increased the spectral exponent in both the frequency bands when compared to vehicle-treatment. Moreover, the CCO activity decreased in the case of CHO and IDB while increased in the case of LCLA when compared to vehicle-treatment. These changes may reflect the underlying mechanism of action of the drugs. CHO seems to decrease the CCO activity while increasing the spectral exponent in both the frequency bands, which can be related to impaired recurrent inhibition in the excitatory pyramidal cells^58^. Indeed, distinct inhibitory circuits orchestrate the oscillations in the cortical circuits^59^, which complicates the E-I balance hypothesis for pharmacology where the relative activity of different subtypes of excitatory or inhibitory neurons can be affected differently by the drugs^11^. Here, all the drugs seem to reduce the oscillatory activity in the 10 – 15Hz frequency band when compared to vehicle-treatment condition (Appendix 2, Figure A2(D)). Also, CHO seems to enhance the oscillatory activity between 30 – 35Hz (Appendix 2, Figure A2(E)) while reducing CCO activity. IDB that affects mitochondrial bioenergetics^36^ (see Figure 3(A)) by mediating electron transfer to complex III in the mitochondrial inner membrane^60^ seems to silence all oscillatory activity (Appendix 2, Figure A2(F)) while decreasing CCO activity. Also, LCLA acted on the activities of the mitochondrial complexes^37^ and increased the CCO activity, which may be related to the high-frequency broadband activity between 35 – 45Hz (Appendix 2, Figure A2(G)). Such changes in the high-frequency components in the LFPs may represent aggregate spiking activity of the local fast-spiking inhibitory neuronal population^13^, which may be relevant in addition to the changes in the spectral exponent related to the E-I synaptic events.

When investigating the effects of drugs on cerebral organoids from healthy controls (Appendix 3), we found that CHO significantly reduced the CCO activity in the cerebral organoid at two months when compared to vehicle-treatment condition (Appendix 3, Figure A3.1(A)). Also, CHO did not significantly affect the spectral exponent in the 1 – 20Hz frequency band (Appendix 3, Figure A3.1(B)) but increased the spectral exponent in the 30 – 50Hz frequency band (Appendix 3, Figure A3.1(C)) when compared to the vehicle-treatment condition in the cerebral organoid at 2 months from healthy controls. In general, CHO seems to reduce the CCO activity while increasing the oscillatory activity in the 30 – 50Hz frequency band in cerebral organoids from healthy controls, as also found in the case of CHO treated cerebral organoids from SCZ patients at 34 days (Figure 3(C)). In the case of IDB treatment, cerebral organoid at 34 days from healthy controls had a decreased CCO activity when compared to SCZ cerebral organoid (Appendix 3, Figure A3.2(A)). The effect of IDB on the spectral exponent in the 1 – 20Hz frequency band was found similar (Appendix 3, Figure A3.2(B)); however, the spectral exponent in the 30 – 50Hz frequency band was increased in the case of healthy controls when compared to SCZ cerebral organoid (Appendix 3, Figure A3.2(C)). LCLA treatment decreased the CCO activity in the cerebral organoid at 34 days from healthy controls when compared to SCZ patients (Appendix 3, Figure A3.3(A)). Also, the effect of LCLA on the spectral exponent in the 0 – 20Hz frequency band was found similar (Appendix 3, Figure A3.3(B)); however, the spectral exponent in the 30 – 50Hz frequency band was decreased in the case of healthy controls when compared to SCZ patients (Appendix 3, Figure A3.3(C)). Therefore, the spectral exponent in the 30 – 50Hz frequency band was found to be most responsive to the drug treatment, which can be related to the drug effects on the E-I ratio. Here, IDB and LCLA had opposite effects (IDB increased while LCLA decreased the E-I ratio) in healthy controls when compared to SCZ cerebral organoids at 34 months even though both affected the activities of the mitochondrial complexes and led to a decrease in the CCO activity in healthy controls when compared to SCZ patients.

Limitations of this study include a limited number of organoids evaluated with drug treatment. In this study, we used two healthy control lines and two schizophrenia lines from our prior works^8,38^ to evaluate the feasibility of the proposed ‘Phase Zero’ clinical study platform. We evaluated the CCO activity using broadband (490–900 nm) absorption spectroscopy, although the state of other complexes in the mitochondrial respiratory chain (complexes I-IV) may also be relevant. For example, IDB mediates electron transfer to complex III in the mitochondrial inner membrane, which was observed based on the state of the complex IV since IDB can restore oxygen consumption at complex IV (and ATP production) based on prior work^60^. Also, LCLA treatment has been shown to lead to the recovery of the activity of complex I and IV^37^. Spectrophotometric assays can be used to measure the enzymatic activities of all the complexes I–IV^61^,^62^. Nevertheless, CCO is the terminal enzyme of the mitochondrial respiratory chain, so the activity of CCO can provide a measure of bioenergetics (ATP production). Here, ATP production by mitochondria at the pre-synapse and the post-synapse supplies energy for the synaptic transmission at the dendrites where interactions are in both directions^24^. Although synaptic activity, as well as spikes together, can provide the energy budget^63^; however, we only investigated the synaptic E-I ratio based on spectral exponent. Unidimensional E-I balance hypothesis may provide an incomplete picture, as found from the drug responses in this study, where more dimensions, including neural firing rates and correlations, may be relevant^64^. Nevertheless, most brain energy is used on synaptic transmission^20^, and neuropsychiatric disorders frequently involve deficits in synaptic E-I balance^11^ and synaptic energy supply^20^.

We postulate that a multidimensional approach combining features from electrophysiology and vis-NIR spectroscopy can complement traditional drug design approaches, that are based on reversing molecular deficits, to push the system towards normative parameter space. In the current feasibility study, IDB drug treatment performed the best in decreasing the CCO activity and the spectral exponent in the 1 – 20Hz frequency band when compared to the vehicle-treatment. However, all the drugs increased the spectral exponent in the 30 – 50Hz frequency band when compared to the vehicle-treatment where neural firing rates and correlations may also be relevant^64^. Therefore, it is important to investigate the relation between E-I balance^11^ and synaptic energy supply^20^, where recent developments in time-resolved vibrational spectroscopy and other techniques^65^ can provide a more comprehensive picture of the metabolic coupling with the electrophysiological features of synaptic E-I balance^11^ and the distributions of spike covariances^4^. Furthermore, future studies need to investigate closed-loop non-invasive stimulation^66^ as external inputs^5^ to longitudinaly modulate E-I balance in the cerebral organoid’s functional circuits^67^, which may be necessary to transition to a more in-vivo cortical circuit dynamics^52^.

## Data availability

Data available on request from the authors

## Acknowledgements

The technology development to combine broadband vis/near-infrared spectroscopy and electrophysiology for preclinical and clinical studies is funded by the Community for Global Health Equity Seed Funding, University at Buffalo, Department of Biotechnology (DBT), Government of India, the Bill and Melinda Gates Foundation & IKP Knowledge Park, India. Mancheung Cheung and Mahasweta Bhattacharya were partly funded as interns by the Community for Global Health Equity Seed Funding, University at Buffalo.

## Author contributions

AD conceived the electrophysiology and broadband vis/near-infrared spectroscopy experiments. AD, MB, SSK, and MC conducted the electrophysiology and broadband vis/near-infrared spectroscopy experiments. MS and ES conceived the protocol for organoid development and drug treatment. ML, JS, and ES conducted the organoid development and drug treatment. AD, MB, SSK performed statistical analysis and figure generation. AD wrote the manuscript. All authors reviewed the manuscript.

